# Population-level distribution and putative immunogenicity of cancer neoepitopes

**DOI:** 10.1101/192716

**Authors:** Mary A. Wood, Mayur Paralkar, Mihir P. Paralkar, Austin Nguyen, Adam J. Struck, Kyle Ellrott, Adam Margolin, Abhinav Nellore, Reid F. Thompson

## Abstract

**Background:** Tumor neoantigens are a driver of cancer immunotherapy response; however, current neoantigen prediction tools produce many candidates that require further prioritization for research/clinical applications. Additional filtration criteria and population-level understanding may help to produce refined lists of putative neoantigens. Herein, we show neoepitope immunogenicity is likely related to measures of peptide novelty and report population-level behavior of these and other metrics.

**Methods:** We propose four peptide novelty metrics to refine predicted neoantigenicity: tumor vs. paired normal peptide binding affinity difference, tumor vs. paired normal peptide sequence similarity, tumor vs. closest human peptide sequence similarity, and tumor vs. closest microbial peptide sequence similarity. We apply these metrics to tumor neoepitopes predicted from somatic missense mutations in The Cancer Genome Atlas (TCGA) and a cohort of melanoma patients, as well as to a group of peptides with neoepitope-specific immune response data using an extension of pVAC-Seq [1].

**Results:** We show neoepitope burden varies across TCGA disease sites and HLA alleles, with surprisingly low repetition of neoepitope sequences across patients or neoepitope preferences among sets of HLA alleles. Only 20.3% of predicted neoepitopes across TCGA patients displayed novel binding change based on our binding affinity difference criteria. Similarity of amino acid sequence was typically high between paired tumor-normal epitopes, but in 24.6% of cases, neoepitopes were more similar to other human peptides, or even to bacterial (56.8% of cases) or viral peptides (15.5% of cases), than their paired normal counterparts. Applied to peptides with neoepitope-specific immune response, a linear model incorporating neoepitope binding affinity, protein sequence similarity between neoepitopes and their closest viral peptides, and paired binding affinity difference was able to predict immunogenicity with an AUROC of 0.66.

**Conclusions:** Our proposed neoepitope prioritization criteria emphasize neoepitope novelty and refine patient neoepitope predictions for focus on biologically meaningful candidate neoantigens. We have demonstrated that neoepitopes should be considered not only with respect to their paired normal epitope, but with respect to the entire human proteome, as well as bacterial and viral peptides, with potential implications for neoepitope immunogenicity and personalized vaccines for cancer treatment. We conclude that putative neoantigens are highly variable across individuals as a function of both cancer genetics and personalized HLA repertoire, while the overall behavior of filtration criteria reflects predictable patterns.

## Background

Neoepitopes are novel peptides that correspond to tumor-specific mutations, are presented on the surface of tumor cells, and have the potential to elicit an immune response (denoting a “neoantigen”). When targeted by cytotoxic T-cells, tumor-associated neoantigens may be associated with increased survival among some cancer patients (e.g. melanoma, cholangiocarcinoma) [2,3], and tumor neoepitope burden seems to correlate with patient survival [4,5,6]. Increasingly, immune checkpoint inhibitor therapies have been successful at stimulating anti-tumor immune responses in several cancer types [7]. However, this ability to leverage the immune system against tumors remains predicated on the immune system’s ability to recognize tumor neoepitopes as “non-self” [8]. Increased tumor neoepitope burden is associated with response to immune checkpoint inhibitor therapies [9,10], and recent attempts to treat melanoma with personalized neoantigen vaccines have shown preliminary success [11,12].

Importantly, not all tumor mutations produce neoepitopes. First, the mutation must result in a change in the amino acid sequence of the tumor peptide relative to the normal peptide. The resulting peptide must also be expressed within cancer cells and bind with high affinity to one or more of the patient’s major histocompatibility complexes (MHC) [8], the polyprotein complexes predominantly encoded by the polymorphic Human Leukocyte Antigen (HLA) loci, which are responsible for presenting peptides to the surface of both normal and cancer cells for detection by the immune system in a patient-specific manner [13,14,15]. With little variation, these are the criteria applied by all computational tools for neoepitope prediction from tumor genomic sequencing data, including Epi-Seq [16], EpiToolKit [17], pVAC-Seq [1], INTEGRATE-neo [18], TSNAD [19], MuPeXI [20], and CloudNeo [21].

Our central assertion is that the immunogenicity of a neoepitope is directly related to its novelty: that is, the extent to which it or a closely matching peptide has previously been presented to the immune system. There is emerging evidence that at least four such novelty criteria may be important to incorporate when identifying and filtering candidate neoepitopes:

1. A neoepitope with strong affinity for MHC (<500 nM [22]) may be a more robust neoantigen candidate if the paired normal epitope has a poor affinity for MHC (>500 nM). This concept was already implemented in the CloudNeo tool, but the effects of filtering epitope calls in this fashion were not addressed [21]. A greater difference in MHC binding affinity between tumor and normal epitopes can increase neoepitope immunogenicity, as shown by Duan et al. using a “differential agretopicity index” [16].
2. While a tumor neoepitope may bind differently from its paired normal epitope, decreased peptide-peptide similarity of the pair at non-MHC-anchoring residues (e.g. amino acid positions 2 and 9 for a 9mer peptide [23]) is likely to increase neoepitope immunogenicity, per the criterion of continuity hypothesis proposed by Pradeu and Carosella, which suggests that epitopes discontinuous with those that the immune system normally encounters are more likely to trigger an immune response [24]. In fact, Yadav et al. found that neoepitopes with amino acid changes at solvent-exposed positions elicited strong T-cell responses [25].
3. Though most approaches only consider the paired normal epitope as a counterpart to its neoepitope, the tumor neoepitope may actually be highly similar to other normal peptides; this emphasizes the importance of considering sequence homology of a neoepitope not just to its normal counterpart, but to all normal peptides that the immune system may encounter in the body. This idea has been previously addressed by the tool MuPeXI [20], but only by searching for *exact* sequence matches of neoepitopes to the reference proteome. Others have investigated the importance of neoepitope sequence similarity to known antigen sequences in predicting response to immunotherapy [26].
4. It is important to consider the sequence homology of candidate neoepitopes to bacteria and viruses, as a) immunotherapy response has shown dependence upon commensal bacteria [27,28,29], b) peptides of bacterial and viral pathogens can be cross-reactive with tumor peptides and recognized by the same tumor-specific T cells [30], and c) virus-derived oncoproteins from virus-associated cancers such as head and neck cancer [31], cervical cancer [32], and Merkel cell carcinoma (MCC) [33] have been shown to elicit T-cell responses [34].

Based on the above data and phenomena, we propose to incorporate the following four biologically significant metrics, summarized in Figure 1, as an extension of pVAC-Seq, with potential for use with other neoepitope calling tools:

1. **Tumor vs. paired normal peptide binding affinity difference**: the degree to which the change in predicted MHC binding affinity between tumor and normal epitopes may be immunogenic.
2. **Tumor vs. paired normal peptide sequence similarity**: the similarity between paired tumor and normal epitopes, based on protein sequence similarity measures as computed from a BLOSUM62 matrix [35].
3. **Tumor vs. closest human peptide sequence similarity**: the similarity between the neoepitope and other normal, unrelated human peptides.
4. **Tumor vs. closest microbial peptide sequence similarity**: the similarity between the neoepitope and known bacterial and viral peptides (from commensals or other infectious pathogens).

**Figure 1.**
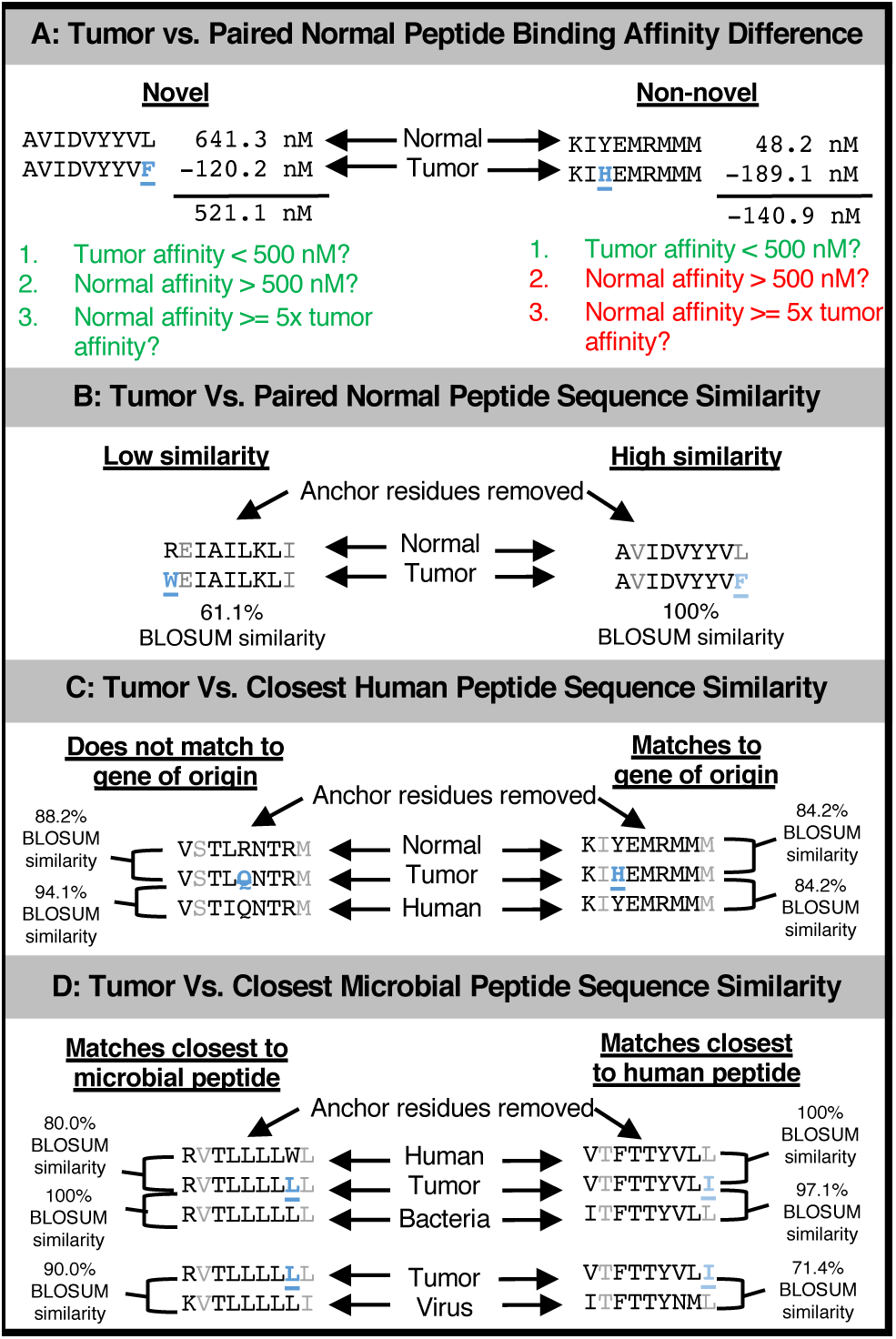
Illustration of proposed neoepitope prioritization metrics. **A**. Tumor vs. paired normal peptide binding affinity difference addresses the difference in MHC Class I binding affinity between the paired tumor and normal epitopes, and a novel binding change occurs when a tumor epitope binds readily to a patient’s HLA allele while its paired normal epitope does not. Examples are shown of a neoepitope which displayed a novel binding change (left) and a neoepitope which did not (right). Mutated residues are shown in blue. **B**. Tumor vs. paired normal peptide sequence similarity addresses the similarity in sequence between the paired tumor-normal epitopes at non-anchor residues based on a BLOSUM62 matrix, normalized by the tumor epitope’s similarity with itself. Examples are shown of a neoepitope with low similarity to its paired normal epitope (left) and a neoepitope with high similarity to its paired normal epitope (right). Anchor residue positions are shown faded, and mutated residues are shown in blue and underlined. **C**. Tumor vs. closest human peptide sequence similarity addresses how similar the neoepitope is to all human proteins based on a blastp search. Examples are shown of a neoepitope which matched to a peptide from a gene other than its gene of origin (left) and a neoepitope which matched to a peptide from its gene of origin (right). Anchor residue positions are shown faded, and mutated residues are shown in blue and underlined. **D**. Tumor vs. closest microbial peptide sequence similarity addresses how similar the neoepitope is to all bacterial and viral proteins based on a blastp search. Examples are shown of a neoepitope that matches closer to a microbial peptide than any human peptide (left) and a neoepitope which matches closer to a human peptide than any microbial peptide (right). Anchor residue positions are shown faded, and mutated residues are shown in blue and underlined.

Herein, we apply these metrics to neoepitopes predicted for somatic mutations identified in a cohort of melanoma patients and across 18 diseases in The Cancer Genome Atlas (TCGA), with the aim of understanding how these metrics influence and stratify neoepitope predictions, both for an individual and at a population level. To our knowledge, our analysis of neoepitope predictions from TCGA represents the broadest study of this kind to date, describing variation across not only a large patient cohort, but across HLA alleles encompassing 99% of the variation in the population at these loci. We also apply our metrics to a small cohort of individual peptides to assess their efficacy of immunogenicity prediction.

## Methods

### pVAC-Seq Analysis of The Cancer Genome Atlas patients

For our analyses, we used somatic mutations identified with MuTect [36] from Mutation Annotation Format (MAF) files for 18 cancer types (see Supplementary Table S1) in TCGA, retrieved using gdc-scan (v1.0.0) [37]. The MAF files were then converted to tumor-normal pair variant call format (VCF) files using the maf2vcf tool in the vcf2maf software package [38], with the GRCh38/hg38 genome build available from the Broad Institute resource bundle [39] used as the reference genome. Because these VCFs still contained data for both tumor and normal samples, they were then manipulated to remove data from the paired normal sample, leaving final, tumor-only VCF files for compatibility with pVAC-Seq, which accepts only single-sample VCFs. Each disease type consisted of a variable number of patients (see Supplementary Figure S1).

We then annotated these VCF files using Variant Effect Predictor (VEP, v88) [40]. VEP was run according to pVAC-Seq’s recommendations [41] with the Downstream and Wildtype VEP plugins [42] used, gene symbols added to output where available, and mutation consequence terms based on Sequence Ontology annotation guidelines [43]. We also used VEP’s GRCh38 annotation cache (rather than querying remotely) for efficiency.

As the TCGA data we obtained did not allow us to calculate patient-specific HLA types, we assumed each tumor could occur in the setting of any HLA allele type, allowing us to explore neoepitope distributions among a broader theoretical population. To do this, we generated a list of HLA alleles to use for subsequent analysis based on allele frequencies originating from the Allele Frequency Net Database [44] and summarized for use in the software POLYSOLVER (v1.0) [45]. The average frequencies across races (Asian, Black, and Caucasian) of alleles for each HLA gene (HLA-A, HLA-B, and HLA-C) were calculated and normalized to sum to 100%. We then selected the top 145 average-frequency HLA alleles for subsequent analysis (see Supplementary Tables S2 – S4), encompassing all HLA alleles among 99% of individuals in the general population.

pVAC-Seq (v4.0.8) was run for each patient and allele combination using 9mer epitopes generated from 17-mer peptides surrounding each missense mutation, and using MHC binding predictions generated by NetMHCpan (v2.8) [46]. For each resulting neoepitope from pVAC-Seq, additional metrics were applied as described below. Note that for the purposes of this study, only epitopes resulting from missense mutations were considered for further analysis, and all peptides were considered to be expressed at equal levels. Note also that neoepitopes from breast cancer (BRCA), cervical cancer (CESC) and melanoma (SKCM) were not assessed for protein sequence similarity against peptides other than their paired normal epitopes.

To assess the degree to which HLA alleles might have overlapping preference for putatively novel binding neoepitopes predicted for mutations across TCGA (see Neoepitope Prioritization Metrics), 1000 random sets of six of the previously described set of HLA alleles (two HLA-A, two HLA-B, and two HLA-C alleles) were chosen using the random.sample function (without replacement) from the random module in Python 2.7.13 [47] (the combinations tested are available in Supplementary Table S5). All unique amino acid sequences of neoepitopes that bound to one or more alleles within each random allele set were counted; separate counts were kept for neoepitopes that bound to one, two, three, four, five, or six of the six alleles (i.e. increasing levels of overlap). The script for randomly sampling allele sets and determining overlap is available in our GitHub repository [48].

For comparison, we assessed recurrence rates among 2,813,809 simulated neoepitopes (9mers) mirroring the size of the TCGA data set. These neoepitopes were drawn randomly from the GRCh38 peptidome, with subsequent introduction of a random single amino acid substitution at a random position along each 9mer. These simulated peptides were labeled by patient and disease site to produce a random set of peptides for each patient equivalent in size to that patient’s predicted neoepitope repertoire. We repeated this process again for a smaller set of 1000 simulated neoepitopes to assess trends in peptide similarity scores. The gene of origin of the random peptide and the gene corresponding to its closest peptide match in the human proteome were retained for protein sequence similarity analysis (see “Tumor vs. closest human peptide sequence similarity” below).

### Analysis of neoepitopes in melanoma patients

We identified patient-specific neoepitopes in whole exome sequencing data from 12 patients selected from a study exploring genomic features of response to immunotherapy in melanoma patients [49]. Reads were aligned against the GRCh37d5 reference genome using the Sanger cgpmap workflow [50]. This workflow uses bwa-mem (v0.7.15-1140) [51] and biobambam2 (v2.0.69) [52] to generate genome coordinate-sorted alignments with duplicates marked. Realignment around indels and base recalibration were performed using Genome Analysis Toolkit (v3.6) [53]. Variants were called using VarScan (v2.3.9) [54] in accordance with the methods outlined in the workflow [50]. VCF files were annotated using VEP (v88) [40] as described above. For all missense single nucleotide variants identified, the tumor and normal protein epitopes of 8, 9, 10, and 11 amino acids in length were produced by reconstructing the nucleotide sequence surrounding the mutation using its coordinates from the VCF file and the CDS in the hg19 gene transfer format file [55], and translating this sequence into amino acids. Each patient’s HLA type was determined from FASTQ files using Optitype (v1.3.1) [56], and the binding affinity of all predicted tumor and normal epitopes was predicted with NetMHCpan (v2.8) [46] for each epitope and patient-specific HLA allele combination. Additional prioritization metrics were applied as described below.

### Neoepitope prioritization metrics

All neoepitope novelty metrics are summarized in Figure 1, and scripts for calculating these metrics are available on our GitHub repository [48].

#### Tumor vs. paired normal binding affinity difference

The difference in MHC binding affinity was calculated using NetMHCpan (described above) as the tumor peptide binding affinity subtracted from the normal peptide binding affinity. A novel binding change was defined as a case in which the tumor neoepitope had an MHC binding affinity below 500 nM (tighter association), while the corresponding normal epitope had at least a 5-fold weaker MHC binding affinity (minimum 500 nM). Note that if an unrelated human peptide was found to be similar to the neoepitope (see “Tumor vs. closest human peptide sequence similarity” below), binding affinity difference was also calculated using this peptide’s sequence.

#### Tumor vs. paired normal peptide sequence similarity

Using a BLOSUM62 matrix, the amino acids at each position along the paired tumor and normal epitopes were given an aggregate similarity score, with higher scores indicating higher similarity. We modified the process described by Henikoff and Henikoff [35] to remove known MHC anchor residues (the second residue and last residue of each 9mer epitope) from scoring in order to remove redundancy with the binding affinity difference metrics, and to place emphasis upon residues that may be more accessible for recognition by T-cells [57]. However, because these scores vary depending on amino acid composition of the proteins tested, we performed a normalization: we divided the similarity score for a neoepitope compared to another peptide by the similarity score of the neoepitope tested against itself to produce percent similarity scores. Note that if an unrelated human, bacterial, or viral peptide was found to be similar to the neoepitope (see “Tumor vs. closest human peptide sequence similarity” and “*Tumor vs. closest microbial peptide sequence similarity”* below), paired sequence similarity was also calculated using this peptide’s sequence instead of the paired normal epitope.

#### Tumor vs. closest human peptide sequence similarity

Using BLAST+ [58], a protein-protein, local, ungapped alignment search of all known human proteins was performed to find the closest matching peptide to each tumor peptide. This was performed using blastp (v2.4.0+) and a peptide database constructed with makeblastdb (v2.4.0+) using Ensembl’s set of all GRCh38 peptides [59]. The BLOSUM62 scoring matrix, ungapped alignments, and an E value cap of 200,000 (to capture blast hits for as many epitopes as possible) were applied, and composition-based statistics were turned off. The top scoring alignment (i.e. lowest E value) with an alignment length of 9 was used as the best match for each neoepitope. If more than one alignment shared the top score, the names of all matching peptides were retained. If a top match was the neoepitope’s normal counterpart, a status of “matching” was assigned to the neoepitope, otherwise a status of “nonmatching” was assigned.

#### Tumor vs. closest microbial peptide sequence similarity

Using BLAST+ [58], a protein-protein, local, ungapped alignment search of all known bacterial and viral peptides was performed to find the closest matching bacterial and viral peptides to each tumor peptide. This was performed using blastp (v2.4.0+) and peptide databases made using makeblastdb (v2.4.0+). The bacterial database was assembled using the National Center for Biotechnology Information (NCBI)’s nonredundant bacterial FASTA releases from RefSeq [60], while the viral database was assembled using NCBI’s nonredundant viral FASTA releases from RefSeq [61]. The BLOSUM62 scoring matrix, ungapped alignments, and an E value cap of 200,000 were applied, and composition-based statistics were turned off. The top scoring alignment (i.e. lowest E value) with an alignment length of 9 was used as the best match for each neoepitope for both the bacterial and viral alignments. If more than one alignment shared the top score, the names of all matching peptides were retained for both the bacterial and viral alignments.

### Features associated with immunogenicity

To assess how well our prioritization metrics reflect a neoepitope’s ability to elicit an immune response, we applied our criteria to predicted neoepitopes from six studies in which peptide-specific immune responses were measured [3,11,20,62,63,64]. For data from all studies, we used only peptides which had complete information regarding the neoepitope and its paired normal peptide, as well as complete data regarding epitope-level immune response, providing a total cohort of 419 peptides. Because only binary immune response data was available from Ott et al. [11] and Bjerregaard et al. [20], we generated binary response data from the Carreno et al. [62] dataset for compatibility: a neoepitope was considered to have elicited an immune response if it had a percent neoantigen-specific T-cell in lymph+/CD8+ gated cells of greater than 10%. Of the seven peptides from the Le et al. [64] dataset that were tested for clonal T cell expansion, the three peptides that demonstrated clonal T-cell expansion were considered to have elicited an immune response, while those that only demonstrated immune reactivity in an ELISpot assay were considered not to have elicited an immune response. Among peptides evaluated in co-culture experiments from Tran et al. [3] and Gros et al. [63], those that were T-cell reactive were considered to have elicited an immune response. We produced a linear model to determine the relationship between our neoepitope novelty criteria and peptide-specific immune response (see “Statistical analysis” below). Using Scikit-learn for Python [65], SVM and Random Forest models were also trained with 10 fold cross validation for comparison.

### Statistical analysis

Statistical analysis was performed using R (v3.3.2) in RStudio. To test the relationship between per-allele neoepitope burden and neoepitope frequency across the TCGA dataset, we obtained the Pearson’s product-moment correlation and associated p-value using a two-sided test. To determine whether a difference in tumor vs. paired normal peptide binding affinity difference exists between epitopes with and without an amino acid change at an anchor position, we applied a Wilcoxon rank sum test. We also used the Wilcox test to compare tumor vs. paired normal peptide sequence similarity scores between epitopes with novel vs. non-novel binding changes, and to compare the difference in tumor vs. paired normal peptide sequence similarity scores for the neoepitope with its paired normal epitope and its closest matching human peptide from BLAST for matching versus non-matching genes. A Welch’s two sample t-test was used to compare the similarity of neoepitopes to bacterial vs. viral peptides. We used the package pROC to obtain AUROC scores and the lm function to determine the relationship between our continuous predictors and observed peptide-specific immune response data. Our analyses are available as an R script on our GitHub repository [48].

## Results

### Neoepitope Frequencies

Consistent with prior analyses of TCGA data [4,67], we demonstrate a varied spectrum of neoepitope burden across diseases, with skin cutaneous melanoma and pheochromocytoma/paraganglioma having the highest and lowest median neoepitope burdens, respectively (see Figure 2A). These differences are likely due to known differences in somatic mutation burden as a function of disease type, as the ratio of neoepitopes to somatic missense mutations per patient was relatively constant across diseases with strong correlation between both metrics (Pearson’s product-moment correlation of 0.99; p < 2.2×10^−16^; see Supplementary Figure S2). Across disease types, an average of 36.8% (sd = 0.7%) of all predicted neoepitopes were from HLA-A alleles, 42.4% (sd = 14.8%) from HLA-B alleles, and 21.0% (sd = 0.7%) from HLA-C alleles.

**Figure 2.**
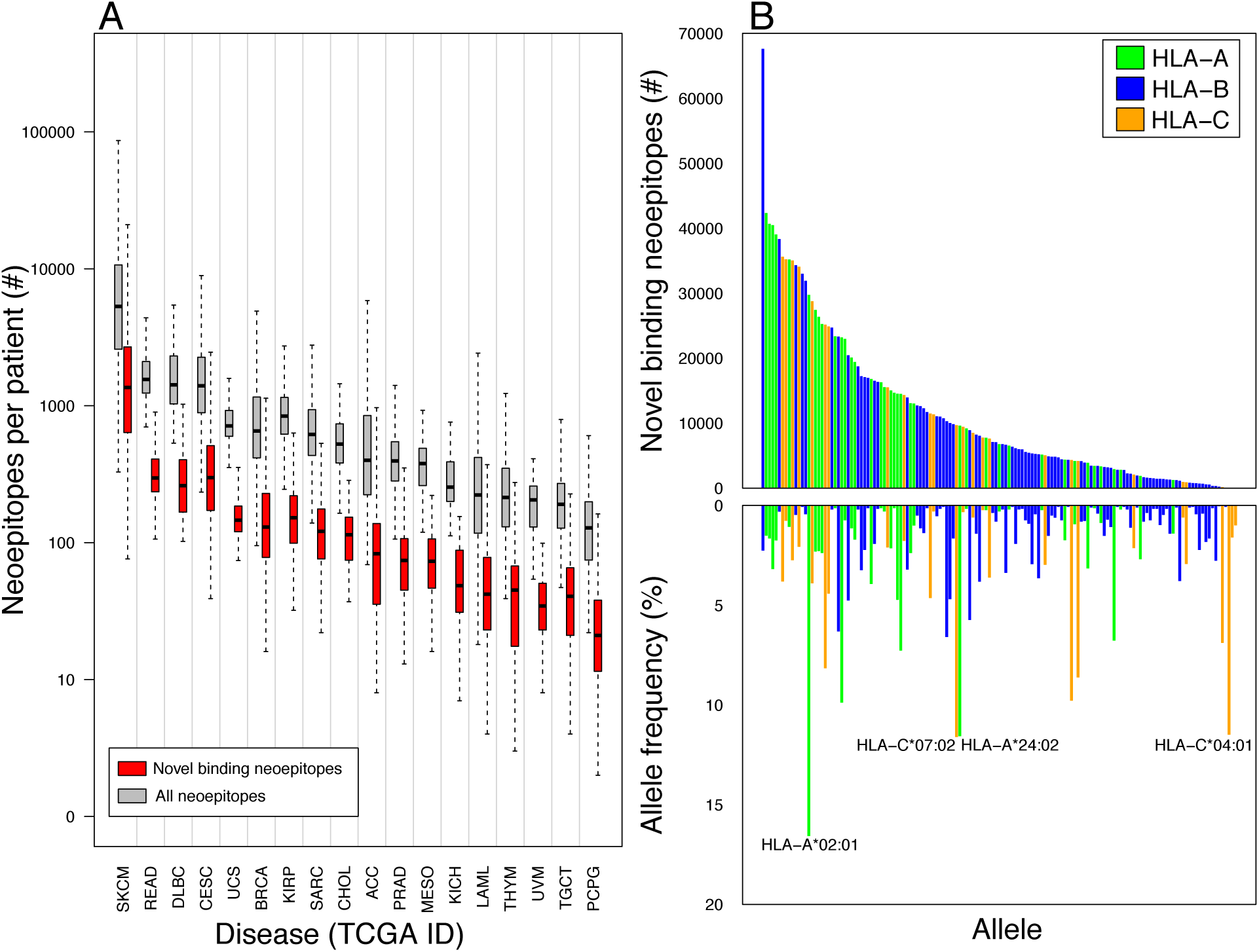
Neoepitope predictions in TCGA across disease sites and HLA alleles. **A**. Number of total and putatively novel-binding predicted neoepitopes in each disease group from TCGA. The total number of neoepitopes (gray) for each patient in each disease group, show in order of decreasing median neoepitope burden, was determined using pVAC-Seq. Novel binding (red) are the subset of neoepitopes which displayed a putatively novel binding change (see Methods). Vertical lines separate TCGA disease sites. Outliers have been removed for clarity. On average, 20.3% of a patient’s neoepitopes were novel-binding. A TCGA disease abbreviation key is available in Supplementary Table S1. **B**. Putatively novel binding predicted neoepitopes across HLA alleles in TCGA. Top: number of neoepitopes with a novel binding change in TCGA for each HLA allele studied, colored green, blue, and yellow according to HLA allele types (A, B, and C, respectively). Bottom: average population frequency for each HLA allele studied, colored as per Top pane. Alleles with >10% frequency in the population are labeled. There was no relationship between allele frequency and number of putatively novel binding neoepitopes (Pearson’s product-moment correlation of 0.1, p=0.1).

We next sought to explore the repertoire of shared neoepitopes across TCGA as a function of HLA subtypes. Among the ten HLA alleles with the greatest number of high-affinity epitopes, there was surprisingly little repetition of epitopes binding to that allele across TCGA (mean 1.06 epitopes repeated); however, each allele had at least one epitope encountered multiple times across patients and diseases (31–168 occurrences). The most frequently repeated neoepitope for HLA-B*15:03, KQMNDARHG, was found most often in breast carcinoma patients. In all cases, this neoepitope originated from a H1047R substitution caused by a single nucleotide variant in the gene *PIK3CA*, a known driver mutation [68]. Two other recurrent epitopes, LSKITEQEK and STRDPLSKI, were identified 168 times in breast, cervical, kidney, and prostate cancers for HLA-A*30:01 and HLA-B*15:17, respectively, with both originating from the same oncogenic mutation in *PIK3CA* (E542K substitutions) [68], and were found exclusively in breast carcinoma, cervical squamous cell carcinoma, and prostate adenocarcinoma patients. There were 5175 other occurrences of *PIK3CA* neoepitopes, as well as 7139 *TP53* (a known cancer driver gene [69]) neoepitopes, and 35872 occurrences of neoepitopes originating from *MUC16*, the gene encoding the CA-125 cancer biomarker [70]. On average, 4.7% of patient epitopes were repeated across patients within their own disease site and 7.9% of patient epitopes were repeated across all of TCGA, a rate significantly higher than that anticipated by random chance alone (see Supplementary Figure S3 and Supplementary Table S6), and likely attributable to common cancer mutations. It is, finally, important to note that the true number of shared neoepitopes among cancers is likely to be smaller due to the random assortment of actual HLA alleles across the population.

We were also interested in the degree to which an overlap of neoepitope preferences existed between the HLA alleles studied, as an epitope that binds strongly to more than one of a patient’s suite of HLA alleles would likely be a better candidate for applications such as peptide vaccines. For 1000 randomly sampled sets of six HLA alleles (see Methods), on average, 3.1% of neoepitopes across TCGA had affinity for at least one of the six alleles in each set, emphasizing the importance of a patient’s unique HLA repertoire in neoepitope presentation. Of these epitopes, the majority (87.5% on average) only had affinity for one allele, and 11.0% on average had affinity for two alleles, but there were some cases where epitopes had affinity for all six alleles (0.0007% of epitopes on average, and up to 0.2% of epitopes for one allele set; see Supplementary Figure S4).

### Tumor vs. paired normal peptide binding affinity difference

We then examined the distribution of paired tumor and normal epitope binding affinities across a cohort of melanoma patients to understand the landscape of differential HLA-specific binding affinities [49] (see Figure 3A). While most mutations did not have a large effect on epitope binding affinity (median 71.1 nM binding affinity difference), we were particularly interested in those mutations that changed the binding affinity significantly in favor of the neoepitope (>5-fold increased affinity, see Figure 3A). We applied these criteria to identify neoepitopes with putatively novel binding changes (see Methods), and noted only a small fraction of qualifying neoepitopes from each patient (0.09% on average, see Figure 3B). This dramatic refinement in neoepitopes led us to consider how these criteria might affect neoepitope distribution across a larger cancer cohort from TCGA (see Supplementary Table S1) and among a broad population of HLA types.

**Figure 3.**
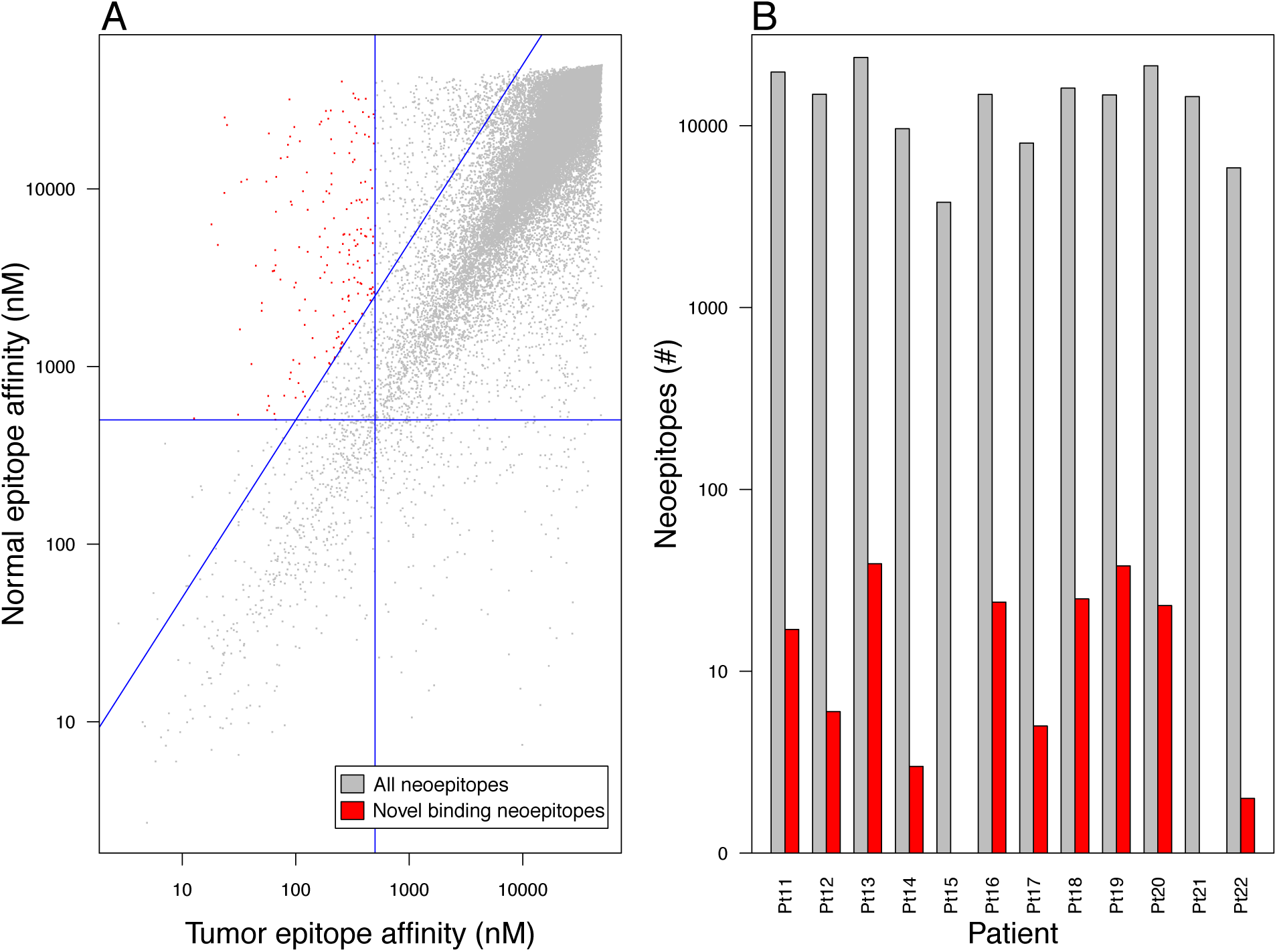
Establishment of putatively novel binding criteria in melanoma patients. **A**. Distribution of paired tumor (x-axis) and normal (y-axis) epitope binding affinities for all neoepitopes analyzed in the Hugo et al. [48] melanoma cohort. The vertical blue line divides epitopes into groups with (left) and without (right) strong tumor epitope binding (MHC affinity <500 nM), while the horizontal blue line divides epitopes into groups with (bottom) and without (top) strong normal epitope binding (MHC affinity <500 nM). The diagonal blue line divides epitopes into groups where the normal epitope binding affinity is at least 5x poorer than the tumor epitope (above) and where the normal epitope binding affinity is less than 5x poorer than the tumor epitope (below). Epitopes colored red are those that have strong tumor epitope binding affinities (<500 nM), weak normal epitope binding affinities (>500 nM), and normal epitope binding affinity at least 5x poorer than tumor epitope binding affinity (constituting a putatively novel binding change). Both axes are shown as log scale. **B**. Number of total (gray) and putatively novel binding (red) neoepitopes for each melanoma patient among the Hugo et al. cohort. Patients 15 and 21 each have a single putatively novel binding neoepitope. Y axis is shown as log scale.

We next assessed novelty of MHC tumor vs. paired normal binding affinity change across TCGA, noting that a minority of neoepitopes (20.3% on average) met this criterion, with a similar distribution across all TCGA cohorts (average proportion per patient of 18.0–24.7%, see Figure 2A). This finding was dependent upon HLA type, with a median 5.6-fold difference in the number of neoepitopes with novel binding changes between the 25^th^ and 75^th^ percentile HLA alleles among diseases (see Figure 2B). All diseases had the greatest number of neoepitopes associated with the allele HLA-B*15:03; however, there was no statistical association between HLA allele frequency in the general population and that allele’s corresponding number of novel binding change neoepitopes (Pearson’s product-moment correlation of 0.1; p = 0.1; see Figure 2B).

Importantly, we note that the mutation of amino acid residues at MHC anchor positions (i.e. the second residue and last residue of each 9mer epitope) tends to result in more dramatic predicted binding affinity differences between the tumor and paired normal peptides compared to mutation of non-anchor residues (median of 2935.1 nM vs. 28.1 nM, respectively; p < 2.2×10^−16^; see Supplementary Figure S5).

### Tumor vs. paired normal peptide sequence similarity

While anchor residue mutations may influence differential peptide binding, anchor residues are anticipated to be more directly engaged with the MHC complex and thus less accessible for T-cell recognition [57]. We therefore sought to investigate the differences in peptide sequence between tumor neoepitopes with mutations in T-cell exposed (i.e. non-anchor) residues and paired normal epitopes. The average protein sequence similarity score (see Methods) between paired epitopes with single nucleotide variants at non-anchor positions across all diseases was 83.5% (ranging from 60.0% to 98.1%, sd = 6.3%). When we assessed these similarity scores in conjunction with our novel binding criteria (see Methods), we observed that neoepitopes which displayed a putatively novel binding change tended to have lower similarity to their paired normal counterpart than those without such a binding affinity change (mean 81.5% vs. 83.7%, respectively; p < 2.2×10^−16^). This level of significance held even when controlling for tumor neoepitope binding affinity (see Supplementary Table S7).

### Tumor vs. closest human peptide sequence similarity

We reasoned that regardless of how similar or dissimilar a neoepitope may be to its paired normal epitope, it may closely mimic a different normal epitope present within the human proteome (see Figure 1C). Comprehensive blastp analysis of all neoepitopes from all disease types generated human proteome matches for more than 99.9% of peptide queries, with an average protein sequence similarity score of 84.3% (sd = 10.7%). The majority of neoepitopes (77.3% on average) mapped most closely to one or more normal peptides from the same gene (see Figure 4A,B). However, 22.7% of neoepitopes matched more closely to one or more unrelated human peptides; in 3.5% of these cases, the unrelated human peptide was an exact match to the tumor neoepitope across all 9 amino acid positions. This phenomenon is likely stochastic in nature, as 24.9% of simulated neoepitopes (see Methods) matched most closely to unrelated human peptides, with an average protein sequence similarity score of 81.9% (sd = 10.6%), significantly different from that of the TCGA neoepitopes (p = 4.927×10^−13^, Welch Two Sample t-test).

**Figure 4.**
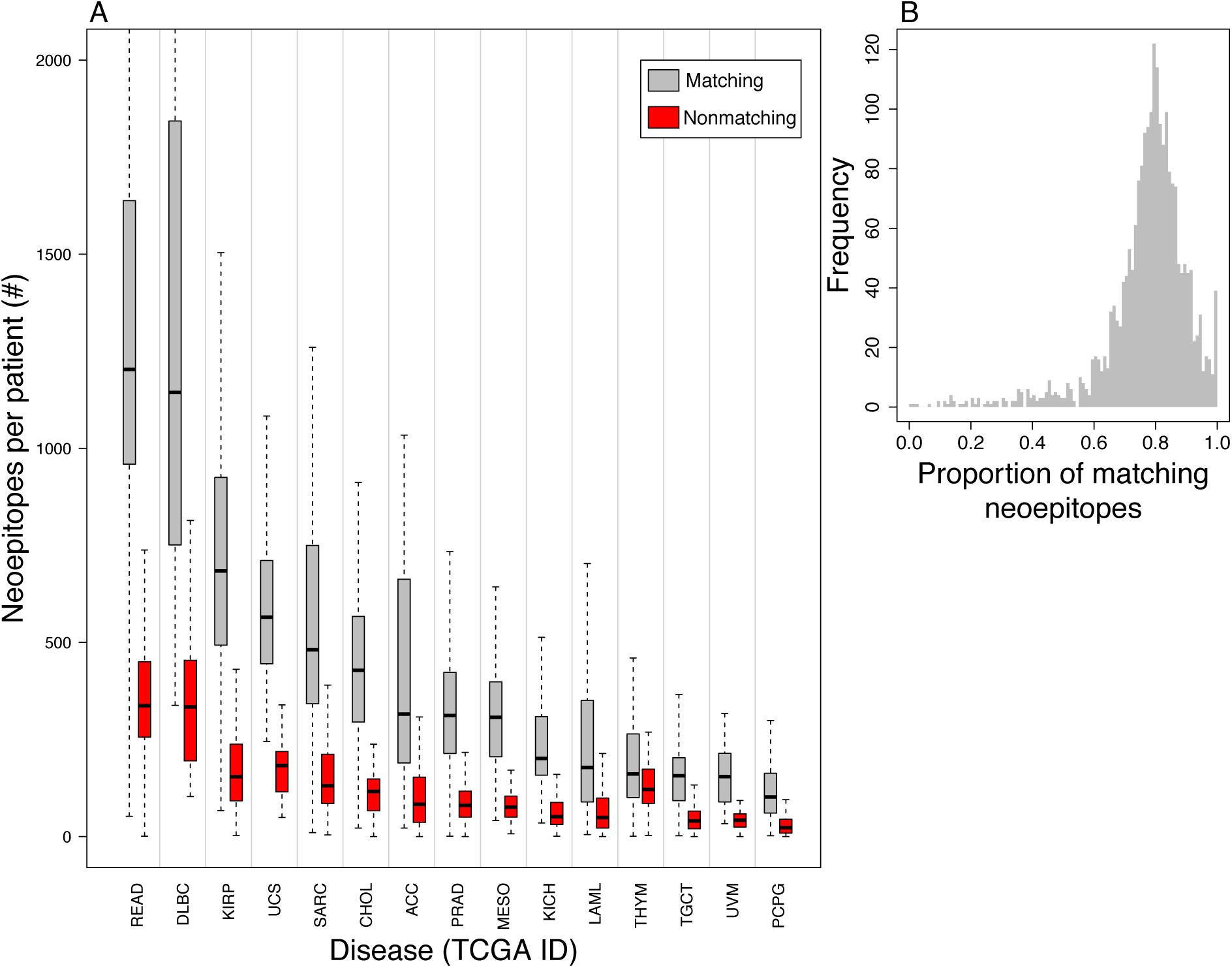
Similarity of neoepitopes to human peptides. **A**. Matching status of top blastp hit(s) to a neoepitope’s gene of origin (see Methods) for patients across TCGA disease groups (gray=matching, red = non-matching). Vertical lines separate TCGA disease sites. Outliers have been removed for clarity. A disease abbreviation key is available in Supplementary Table S1. **B**. Distribution of the proportion of neoepitopes that matched to the gene of origin for each patient’s neoepitopes. On average, 77.3% of neoepitopes for each patient had a top blastp hit that matched their gene of origin.

### Tumor vs. closest microbial peptide sequence similarity

Next, we assessed neoepitope sequence homology with peptides from pathogenic and commensal microorganisms. Almost all neoepitopes were found to have at least one matching bacterial or viral peptide by blastp (87.6% and >99.9%, respectively). Overall, tumor neoepitopes were more similar to bacterial peptides compared to viral peptides (mean percent peptide sequence similarity score of 91.4% (sd = 6.6%) and 76.7% (sd = 9.1%), respectively, (p < 2.2×10^−16^). Interestingly, in 56.9% of cases where a neoepitope had a bacterial blastp hit, the bacterial peptide was more similar to the neoepitope than either its normal counterpart or its most similar normal protein as determined by blastp; this was only true for 15.8% of the viral peptide matches for neoepitopes (see Figure 1D for example). More strikingly, when considering protein sequence similarity scores across all residues, 59.6% of neoepitopes with bacterial blastp hits had higher similarity to these peptides than to either of their normal peptide matches; only 5.8% of viral epitopes showed this phenomenon. This was true despite the fact that neoepitopes had significantly more mismatches in sequence with bacterial peptides than they did with either their paired normal epitopes (mean 1.5 vs 1; p < 2.2×10^−16^) or their closest matching peptides from blastp (mean 1.5 vs 1.4; p < 2.2×10^−16^). However, in terms of total amino acid length, the bacterial peptide data base was 577.5 times larger than the human peptide database, and the viral peptide data base was only 2.0 times larger, so these phenomena may be in part reflective of these differences.

Supplemental Figure S6 shows the distribution of protein percent similarity scores for bacterial and viral hits for each TCGA disease site and for 1000 randomly simulated neoepitopes (see Methods) across non-anchor residues. Predicted TCGA neoepitopes were significantly more similar than random peptides to their closest matching bacterial peptide (mean 91.4% vs. 82.3%, p < 2.2×10^−16^, Welch Two Sample t-test) and their closest matching viral peptide (mean 76.7% vs. 58.7%, p < 2.2×10^−16^, Welch Two Sample t-test), indicating that this phenomenon of sequence similarity to microorganism peptides may be specific to cancer neoepitopes. We determined the top 10 most frequently occurring bacterial genera in cases where a bacterial peptide was a closer match to a neoepitope than either of its human peptide counterparts (see Figure 5), which includes, of particular interest, frequently pathogenic genera such as *Clostridium* [71], *Mycobacterium* [72,73], and *Vibrio* [74], and the frequently commensal genus *Lactobacillus* [75].

**Figure 5.**
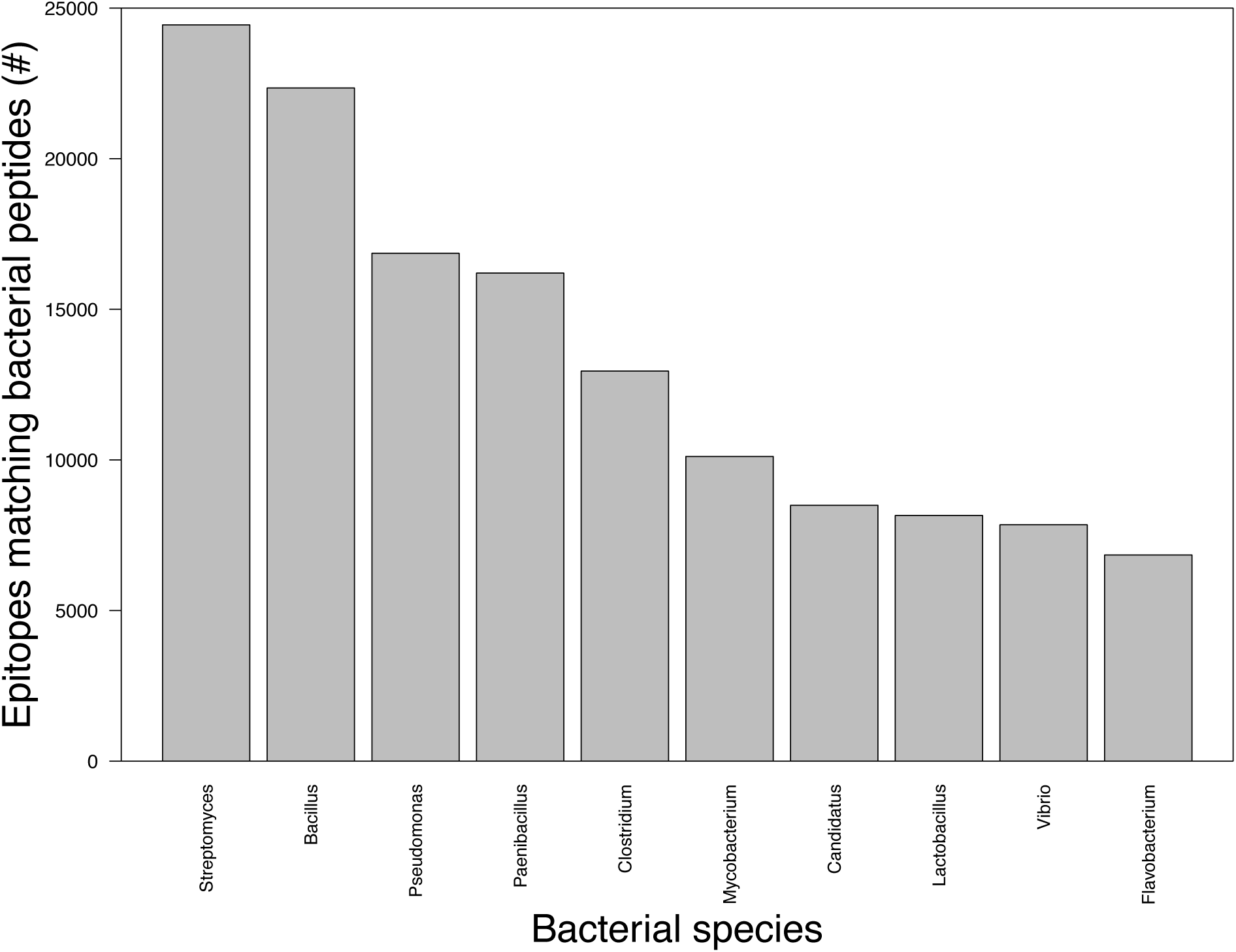
Species of origin of bacterial peptide matches to neoepitopes. Top 10 most frequent bacterial genera with peptides matching more closely to a neoepitope than either its paired normal epitope or its top blastp hit are shown.

### Features associated with immunogenicity

Finally, we applied our criteria to a cohort of neoepitopes with paired immune response data. Applying any single criterion to predict immune response to a neoepitope in a linear model did not lead to significant prediction in any case, except for the percent protein sequence similarity between a neoepitope and its closest viral peptide (p = 0.046; see Table 1, Figure 6A-F). We also observed how well immune response was predicted by neoepitope binding affinity, paired normal epitope binding affinity, and the number of mismatches in amino acid sequence between the neoepitope and its paired normal epitope predicted response. Only the number of mismatches was alone able to predict neoepitope immunogenicity, favoring those neoepitopes with multiple amino acid changes (p = 0.03; see Table 1). Our putatively novel binding change criteria alone was able to predict true immunogenicity with an AUROC of 0.53 (see Supplementary Figure S7). A linear model incorporating 1) neoepitope binding affinity, 2) putatively novel binding change status of the neoepitope, 3) binding affinity difference between the neoepitope and both its paired normal epitope and 4) its closest BLAST peptide match, 5) number of amino acid sequence mismatches between the neoepitope and its paired normal epitope, and 6) percent protein sequence similarity between the neoepitope and its paired normal epitope, 7) its closest human peptide match, 8) its closest bacterial peptide match, and 9) its closest viral peptide match was able to significantly predict immune response to neoepitopes (p = 0.02). However, only three individual predictors contributed significantly to the model: neoepitope binding affinity (p = 0.003), percent protein sequence similarity of the neoepitope to its closest viral peptide match (p = 0.048), and binding affinity difference between the neoepitope and its closest human peptide match (p = 0.002). The contribution of the number of amino acid sequence mismatches between the neoepitope and its paired normal epitope approached significance (p = 0.075). A reduced, multiplicative version of our linear model incorporating only these four predictors was able to predict immune response to neoepitopes with greater significance (p = 3.3×10^−6^; AUROC = 0.66; see Figure 6G; mathematical representation available in supplementary materials). For comparison, we also trained SVM and Random Forest models to predict peptide-specific immune response; however, simpler linear model remained the best predictor (see Supplementary Figure S8).

**Figure 6.**
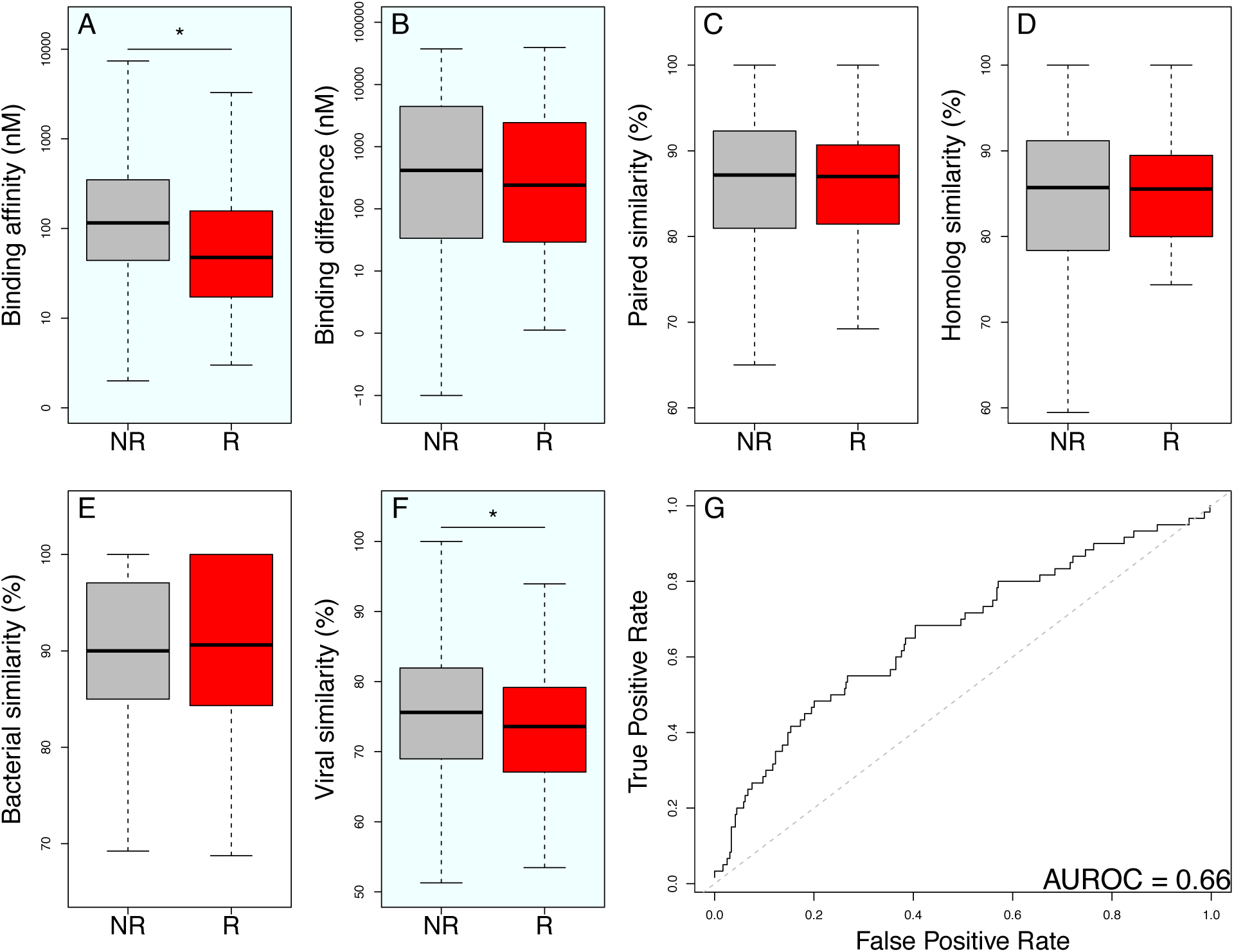
Neoepitope prioritization metric scores and linear modeling in the cohort of peptides with neoepitope-specific immune response data. Score distributions are shown for peptides with (red) and without (gray) neoepitope-specific immune response. Panes in light blue indicate metrics which are included in the model shown in part G. R = “response”, NR = “nonresponse”. **A**. Neoepitope binding affinity, * p < 0.001 in a Wilcoxon rank sum test. **B**. Difference in binding affinity between the neoepitope and its paired normal epitope (normal affinity – tumor affinity). **C**. Percent protein sequence similarity (see Methods) between the neoepitope and its paired normal epitope. **D**. Percent protein sequence similarity between the neoepitope and its closest human peptide. **E**. Percent protein sequence similarity between the neoepitope and its closest bacterial peptide. **F**. Percent protein sequence similarity between the neoepitope and its closest viral peptide. **G**. ROC curve for prediction of immunogenicity from prioritization criteria. A linear model incorporating neoepitope binding affinity, protein sequence similarity between neoepitopes and their closest viral peptides, and difference in binding affinity between neoepitopes and their closest human peptides was used to predict immune response. Within a limited cohort of 419 peptides with immune response data, our model was able to predict peptide immunogenicity with an AUROC of 0.66. Dashed gray line represents the line y=x for comparison.

**Table 1.** Significance of prioritization metrics in predicting immune response. Based on a linear model, each prioritization metric, along with tumor and paired normal epitope binding affinities and the number of sequence mismatches between neoepitopes and paired normal epitopes, were tested for the ability to predict immune response to a predicted neoepitope.

| Predictor of immune response | Adjusted R<sup>2</sup> | Significance (p value) |
| --- | --- | --- |
| Neoepitope binding affinity | -0.002 | 0.7 |
| Paired normal epitope binding affinity | -0.002 | 0.7 |
| Difference in binding affinity between neoepitope and paired normal epitope | -0.002 | 0.7 |
| Difference in binding affinity between neoepitope and closest human protein | -0.002 | 0.3 |
| Number of mismatches between neoepitope and paired normal epitope | 0.01 | 0.03 |
| Percent protein sequence similarity between neoepitope and paired normal epitope | 0.004 | 0.09 |
| Percent protein sequence similarity between neoepitope and closest human protein | -0.001 | 0.5 |
| Percent protein sequence similarity between neoepitope and closest bacterial protein | -0.002 | 0.8 |
| Percent protein sequence similarity between neoepitope and closest viral protein | 0.007 | 0.046 |

## Discussion

In this study, we have explored the frequency and distribution of neoepitopes among patients across diverse TCGA disease types using a broad set of HLA alleles. We have further described and evaluated multiple neoepitope prioritization criteria that can significantly refine patient neoepitope predictions to enrich for biologically meaningful candidate neoantigens. In particular, we proposed four metrics that emphasize neoepitope novelty: tumor vs. paired normal peptide binding affinity difference, tumor vs. paired normal peptide sequence similarity, tumor vs. closest human peptide sequence similarity, and tumor vs. closest microbial peptide sequence similarity. By applying these metrics to predicted neoepitopes from 572,170 combinations of patients and HLA alleles, we estimated the behavior of these metrics in the general cancer population. We have shown that tumor-normal MHC binding affinity differences and non-anchor peptide sequence similarity are independent metrics, and can be used to dramatically refine a list of neoepitopes for further consideration. Applying our criteria to a cohort of neoepitopes with paired immune response data reaffirmed the importance of neoepitope MHC binding affinity in eliciting an immune response. We also demonstrated the significance of novel MHC binding of neoepitopes relative to human proteins, and the degree of sequence similarity of neoepitopes to viral peptide sequences.

To our knowledge, this is the first study to analyze neoepitope predictions broadly across the population by investigating candidate neoantigens among a large cohort of patients within TCGA and across an extensive set of HLA alleles encompassing 99% of the variation in the population at each HLA locus. The approach detailed here also represents the first systematic comparison of neoepitopes to unrelated peptides, demonstrating that a neoepitope may be more similar to other human, commensal, or pathogenic peptides than its paired normal epitope. This approach also builds upon pVAC-Seq in several significant ways, and could in theory be applied as a post-processing step in any neoantigen prediction pipeline. Although the peptide immune response data we analyzed was limited in size and scope, it represents the largest such cohort published to date. We expect further refinement of neoepitope prioritization with additional data and emerging biological insights.

Our study also has several limitations which must be considered when interpreting these results or applying a similar approach prospectively. For simplicity’s sake, we did not consider expression levels or variant allele frequencies of the neoepitopes analyzed, which are important and well-established criteria for prioritizing predicted neoepitopes which are robustly present in the tumor of interest. We also did not enrich *a priori* the subset of microorganisms most closely associated with human health and disease, thus broadening the peptide search space to include likely uninformative sequences (e.g. non-human viruses) while excluding some potentially relevant species (e.g. yeasts). Additionally, as we did not have HLA typing information for the TCGA cohort, we were unable to explicitly address combinatorial overlap of epitope preference for HLA alleles on a per-patient basis. Lastly, this analysis only considers single nucleotide missense mutations.

In the future, we aim to include more complex variants such as small insertions and deletions, as these have the potential to produce highly novel, immunogenic neoantigens [67] and are currently omitted from consideration by all but a few neoepitope prediction tools (e.g. MuPeXI [20] and TSNAD [19]). Further, we believe that the incorporation of tumor neoepitope sequence similarity into studies of the microbiome in cancer patients could be important for better understanding patient response to immunotherapy treatment. Incorporating the criteria proposed here into the development of neoepitope vaccines may help refine the vaccine production process and potentially improve the success of cancer treatments.

## Conclusions

In our exploration of neoepitopes across broad populations and sets of HLA alleles, we have evaluated multiple neoepitope prioritization criteria that emphasize peptide novelty, concluding that neoepitopes should be considered not only with respect to their paired normal epitope, but with respect to the entire human proteome, as well as bacterial and viral peptides, with potential implications for neoepitope immunogenicity and personalized vaccines for cancer treatment. We further conclude that the sequences of putative neoantigens are highly variable across individuals as a function of both cancer genetics and personalized HLA repertoire, while the overall behavior of filtration criteria reflects more predictable patterns.

## Additional Files

### Additional File 1

Supplementary Information. Contains Supplementary Figures S1-S8 and Supplementary Tables S1-S4,S6-S7, and the mathematical representation of our linear model predicting neoepitope-specific immune response.

### Additional File 2

Supplementary Table S5. Contains allele sets tested and associated epitope counts from analysis on overlap of epitope preference among HLA alleles.

## Abbreviations

AUROC: Area Under the Receiver Operating Characteristic Curve

DAI: Differential Agretopicity Index

HLA: Human Leukocyte Antigen

MAF: Mutation Annotation Format

MCC: Merkel Cell Carcinoma

MHC: Major Histocompatibility Complex

NCBI: National Center for Biotechnology Information

ROC: Receiver Operating Characteristic

TCGA: The Cancer Genome Atlas

VCF: Variant Call Format

VEP: Variant Effect Predictor

pVAC-Seq: personalized Variant Antigens by Cancer Sequencing

## Declarations

### Acknowledgments

The results shown here are based upon data generated by the TCGA Research Network: http://cancergenome.nih.gov/.

### Availability of data and materials

The sequence data used for our melanoma cohort as published by Hugo et al. [49] are available from the Sequence Read Archive [75] under the accession numbers SRA: SRP067938 and SRA: SRP090294. TCGA variant call sets are available from the TCGA data portal [76]. The peptide and paired immune response data sets used in our cohort to identify features associated with immunogenicity is available within the published articles from which they originated and their associated supplementary materials [3,11,20,62,63,64]. The datasets produced for our analyses and supporting the conclusions of this article are available upon request.

### Authors’ contributions

MAW – project design, data analysis, data interpretation, manuscript preparation and review, MP – data analysis, manuscript review, MPP – data analysis, manuscript review, AN – data analysis, manuscript review, AJS – data analysis, manuscript review, KE – data analysis, manuscript review, AM – project design, manuscript review, AN – data analysis, manuscript review, RFT – project conceptualization and design, data analysis and interpretation, manuscript preparation and review.

### Consent for publication

Not applicable.

### Ethics approval and consent to participate

Not applicable.

