## Supplementary Materials for "Population-level distribution and putative immunogenicity of cancer neoepitopes"

### Supplementary Material

The mathematical representation of our multiplicative linear model predicting neoepitope specific immune response, as described in the "Features associated with immunogenicity" subsection of our results section, is as follows:

$$\mathbf{Y} = -6.6 + 9.4\mathbf{A} + 8.6\mathbf{V} + 9.3\mathbf{B} + 7.0\mathbf{M} - 12.3\mathbf{AV} + 1.01\text{x}10^{-5}\mathbf{AB} - 12.3\mathbf{VB} - 9.4\mathbf{TM} - 9.1\mathbf{VM} - 9.3\mathbf{BM} - 1.2\text{x}10^{-5}\mathbf{AVB} + 12.4\mathbf{AVM} - 1.0\text{x}10^{-5}\mathbf{ABM} + 12.3\mathbf{VBM} + 1.2\text{x}10^{-5}\mathbf{AVBM}$$

where **Y** is the neoepitope specific immune response, **A** is the HLA binding affinity of the neoepitope, **V** is the peptide sequence similarity of the neoepitope to its closest-matching viral peptide, **B** is the HLA binding affinity difference between the neoepitope and its closest-matching human peptide, and **M** is the number of amino acid mismatches between the neoepitope and its paired normal epitope.

**Supplemental Figure S1.** Total number of patients in each TCGA disease group. A disease abbreviation key is available in Supplementary Table S1.

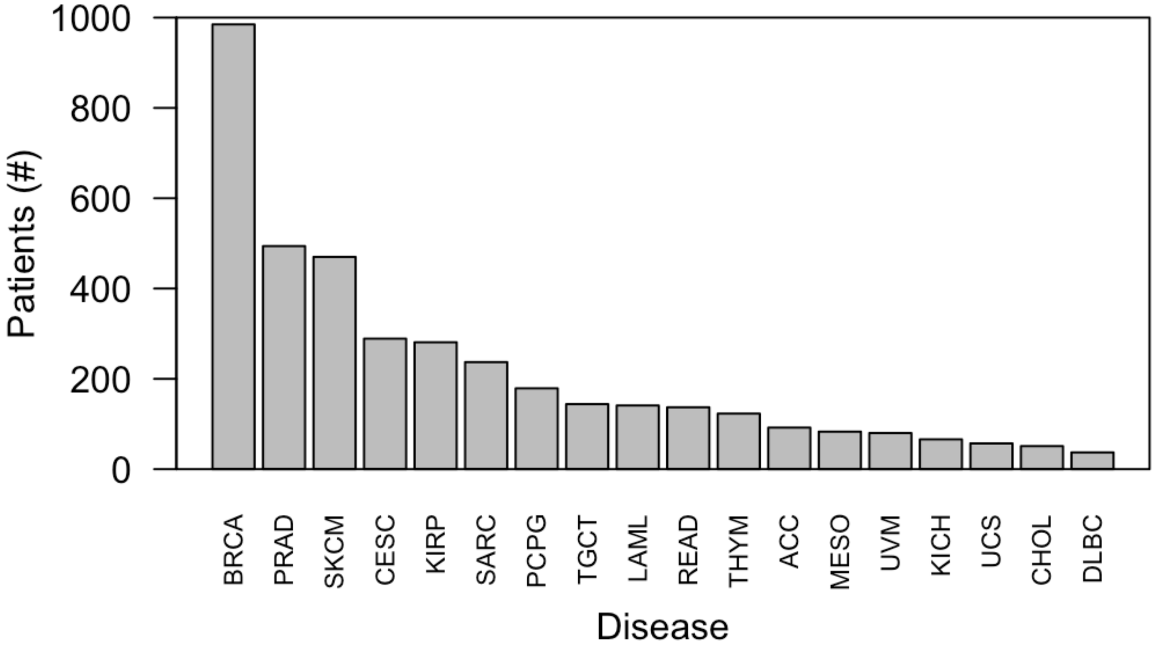

**Supplemental Figure S2.** Ratio of neoepitopes to somatic missense mutations per patient across disease sites. The total number of neoepitopes predicted across all 145 HLA alleles for each patient was divided by the total number of somatic missense mutations identified for each patient to calculate the ratio.

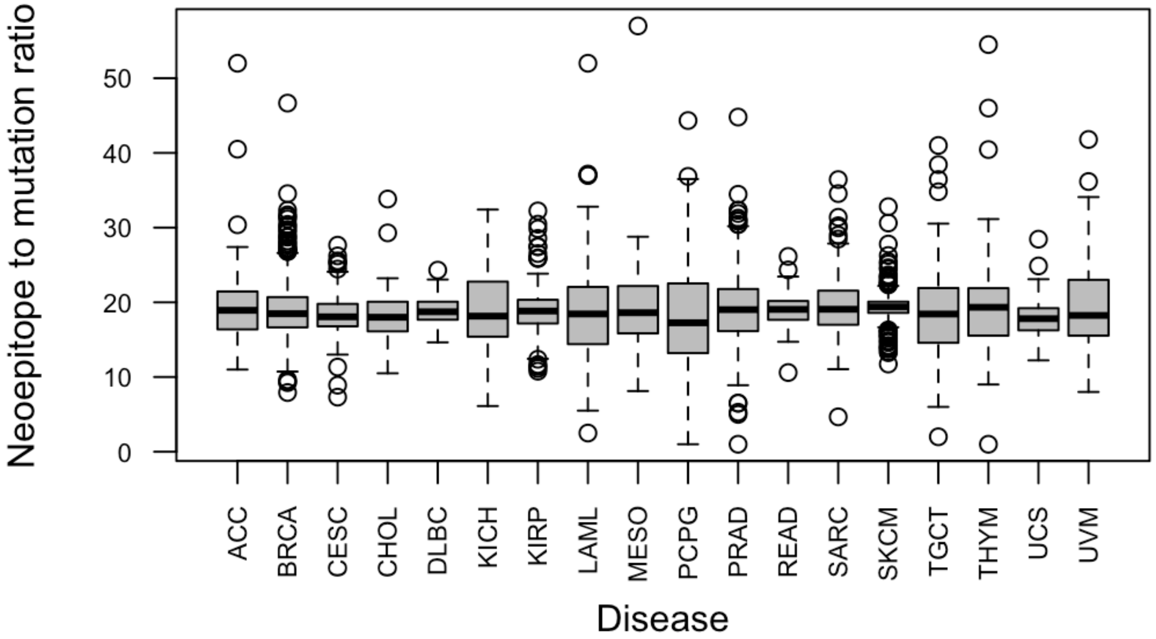

**Supplemental Figure S3.** Distributions of repeated peptide sequences for TCGA epitopes and randomly simulated epitopes (see Methods). **A**, percentage of epitope sequences repeated per patient across patients within the same TCGA disease site compared to the percentage of epitope sequences repeated for the same number of random peptides across TCGA. **B**, percentage of epitope sequences repeated per patient across all patients in TCGA compared to the percentage of epitope sequences repeated for the same number of random peptides across TCGA.

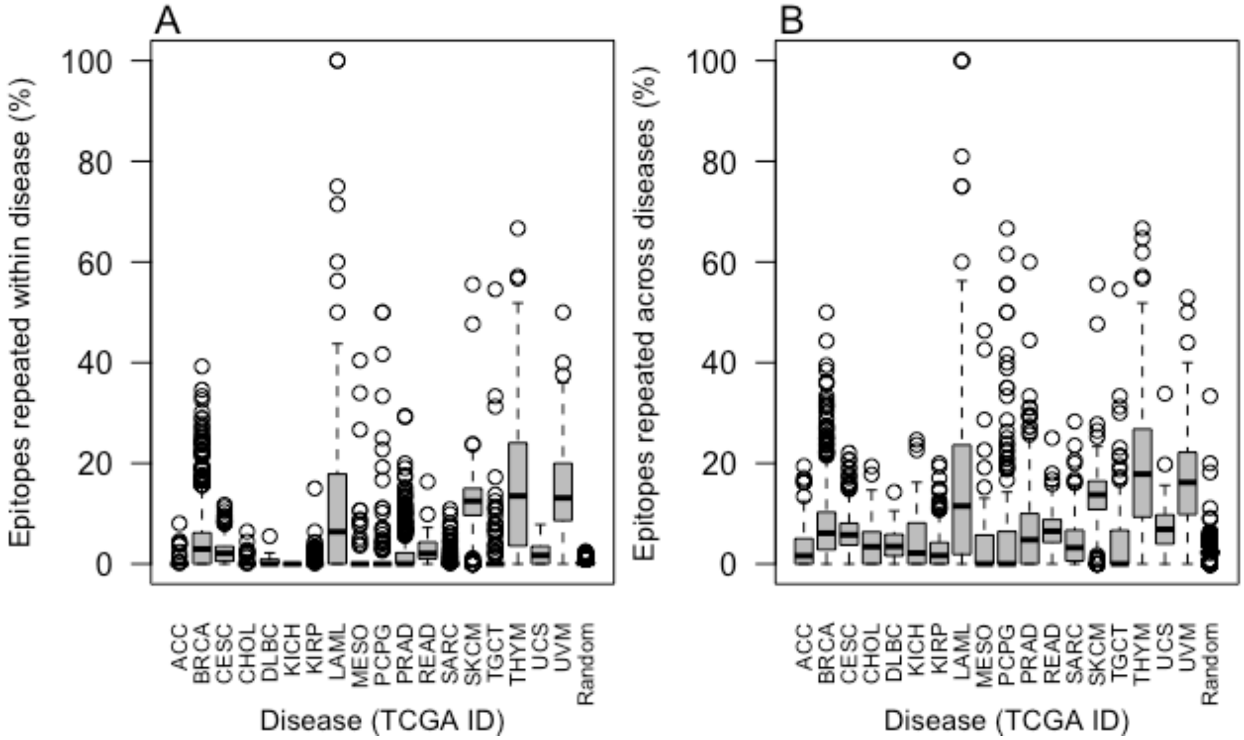

**Supplemental Figure S4.** Overlap of epitope preference rare across HLA alleles. Using 1000 randomly sampled sets of six HLA alleles (2 HLA-A, 2 HLA-B, 2 HLA-C; see Methods), the instances of shared neoepitope preference were assessed for all neoepitopes predicted across TCGA patients. In most cases (87.898% on average), only one allele from each group bound to a neoepitope, but in some cases (0.00433% on average), all six alleles had a strong binding affinity for the neoepitope. The x axis represents the number of alleles (among a set of six) able to present a given neoepitope. The y axis depicts the number (%) of all neoepitopes with affinity for at least one allele in the set presented by the corresponding number of alleles along the x axis, with the distribution representing a summary across 1000 randomly sampled sets of six HLA alleles.

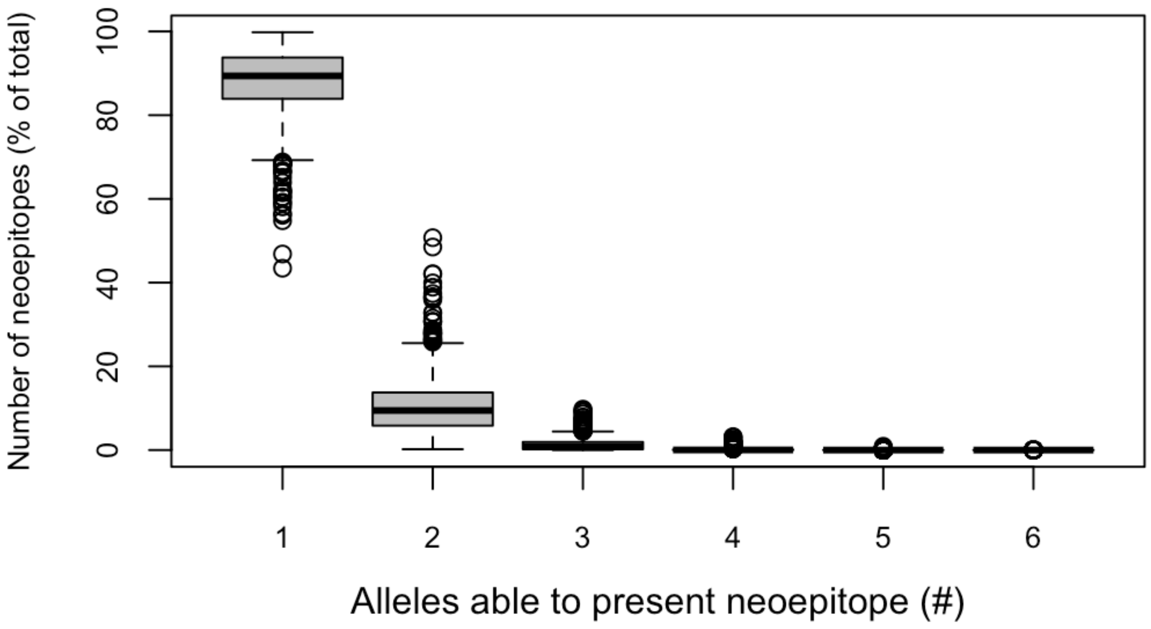

**Supplemental Figure S5.** Binding affinity difference between paired tumor and normal epitopes in TCGA. Binding affinity difference between paired tumor and normal epitopes was higher for neoepitopes with mutations at anchor residues than those with mutations at non-anchor residues (median 2935.1 nM vs. 28.1 nM, respectively; $^*p < 2.2 \times 10^{-16}$).

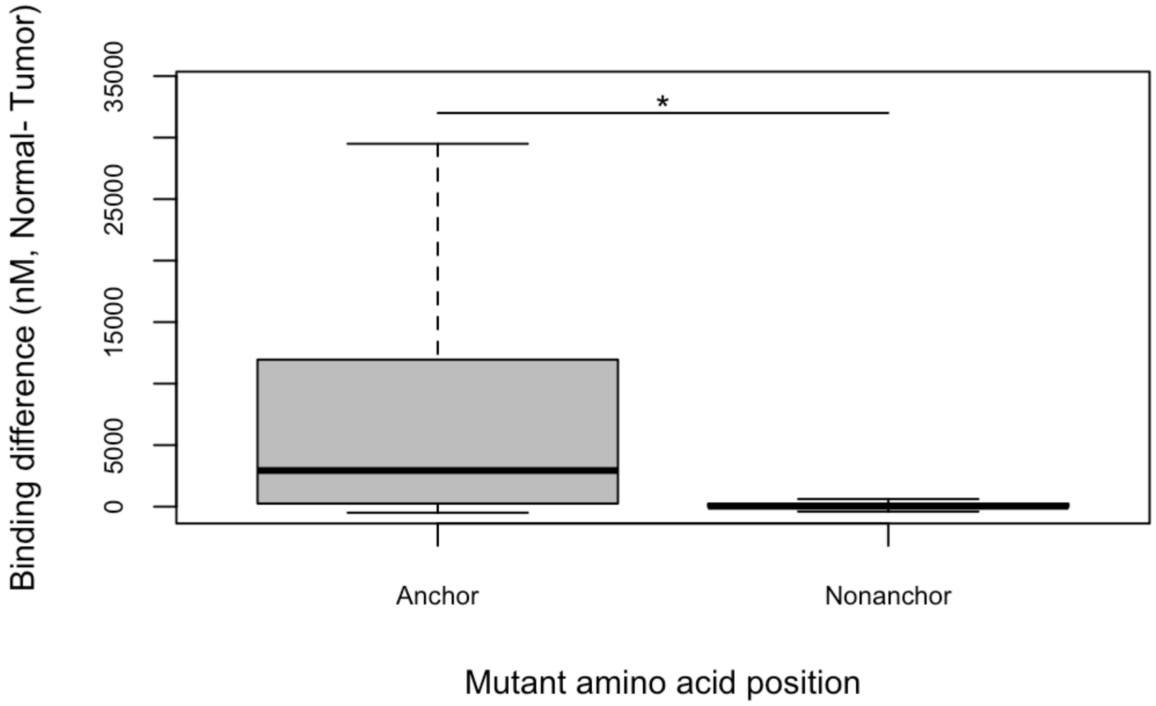

**Supplemental Figure S6.** Tumor vs. closest microbial peptide sequence similarity of predicted TCGA neoepitopes and randomly simulated peptides. Percent paired sequence similarity (see Methods) of neoepitopes with top bacterial peptide (gray) and viral peptide (white) blastp hits across TCGA disease groups and among 1000 randomly simulated neoepitopes (see Methods) is shown. All TCGA neoepitopes matched with some degree of similarity (76.7% on average) to a viral peptide, while 96.4% of neoepitopes matched with some degree of similarity (91.6% on average) to a bacterial peptide. Vertical lines separate TCGA disease sites. A TCGA disease abbreviation key is available in Supplementary Table S1.

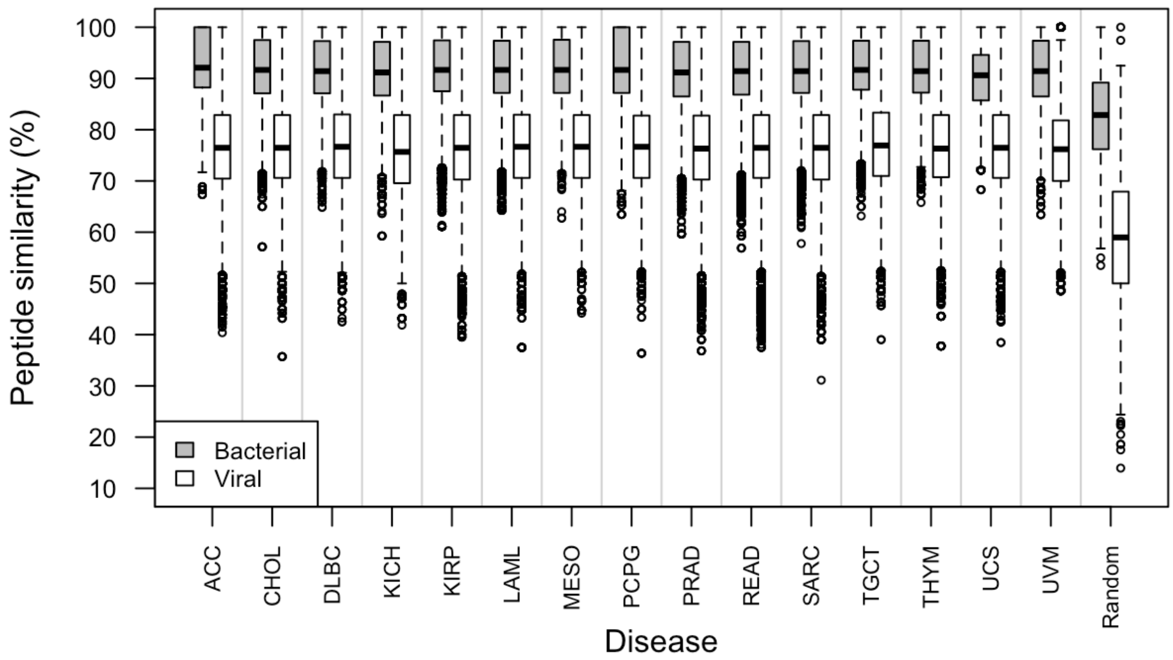

**Supplemental Figure S7.** ROC curve for prediction of immunogenicity from putatively novel binding status. Within a limited cohort of 419 peptides with paired immune response data, our immunogenic binding criteria was able to predict peptide immunogenicity with an AUC of 0.528.

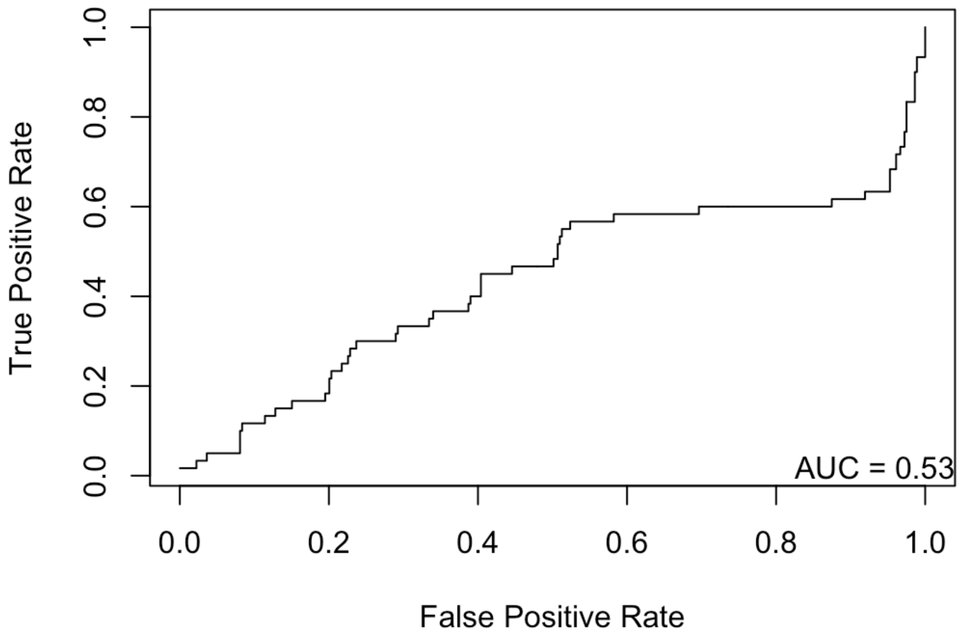

**Supplementary Figure S8.** ROC curves for the prediction of peptide immunogenicity of peptides using Random Forest, SVM, or multiplicative linear models. Random Forest and SVM models were trained with 10-fold cross validation. All models predict neoepitope-specific immune response from 1) neoepitope binding affinity, 2) paired normal binding affinity, 3) difference in binding affinity between the neoepitope and its paired normal epitope, 4) number of mismatches in amino acid sequence between the neoepitope and its paired normal epitope, 5) difference in binding affinity between the neoepitope and its closest matching human peptide, 6) percent protein sequence similarity between the neoepitope and its paired normal epitope, 7) percent protein sequence similarity between the neoepitope and its closest matching human peptide, 8) percent protein sequence similarity between the neoepitope and its closest matching bacterial peptide, and 9) percent protein sequence similarity between the neoepitope and its closest matching viral peptide. Gray line represents the line y=x for comparison.

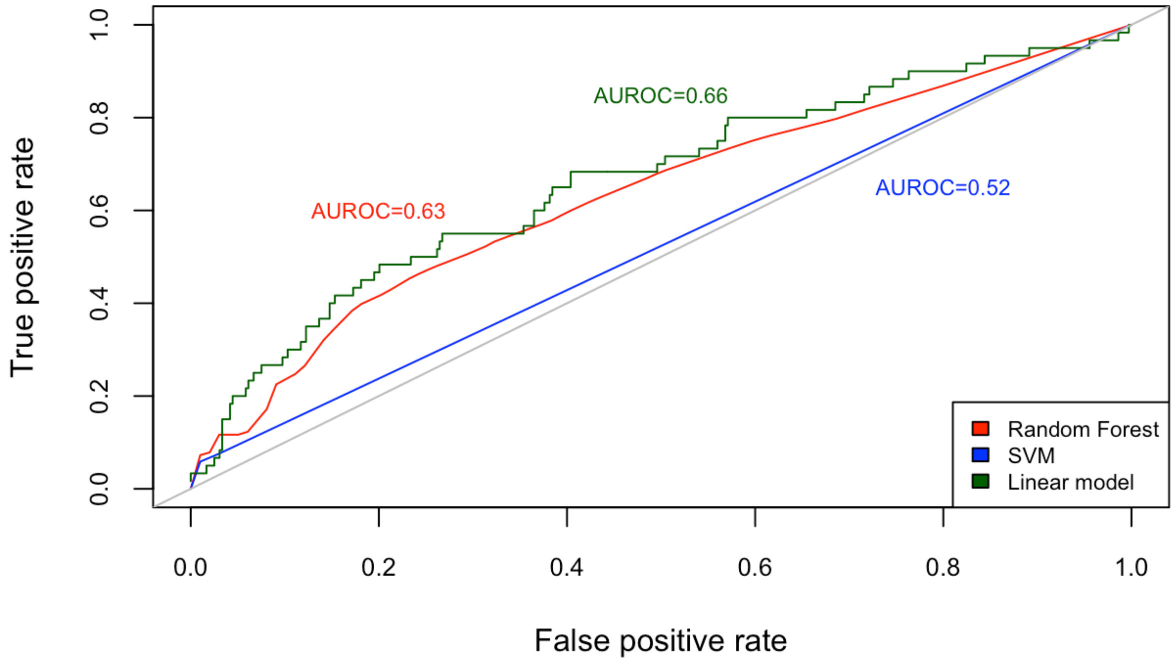

**Supplementary Table S1:** TCGA cancer data sets used. Disease abbreviations and the number of patients used to generate each data set provided.

| TCGA Project Abbreviation | Cancer Type | Number of Cases |
|---|---|---|
| ACC | Adrenocortical Carcinoma | 92 |
| BRCA | Breast Invasive Carcinoma | 985 |
| CESC | Cervical Squamous Cell Carcinoma and Endocervical Adenocarcinoma | 289 |
| CHOL | Cholangiocarcinoma | 51 |
| DLBC | Lymphoid Neoplasm Diffuse Large B-cell Lymphoma | 37 |
| KICH | Kidney Chromophobe | 66 |
| KIRP | Kidney Renal Papillary Cell Carcinoma | 281 |
| LAML | Acute Myeloid Leukemia | 141 |
| MESO | Mesothelioma | 83 |
| PCPG | Pheochromocytoma and Paraganglioma | 179 |
| PRAD | Prostate Adenocarcinoma | 494 |
| READ | Rectum Adenocarcinoma | 137 |
| SARC | Sarcoma | 237 |
| SKCM | Skin Cutaneous Melanoma | 470 |
| TGCT | Testicular Germ Cell Tumors | 144 |
| THYM | Thymoma | 123 |
| UCS | Uterine Carcinosarcoma | 57 |
| UVM | Uveal Melanoma | 80 |

**Supplementary Table S2:** Top frequency HLA-A alleles (n=38) used for neoepitope prediction from TCGA somatic mutations. Average allele frequencies in the general population are based on those frequencies originating from the Allele Frequency Net Database [1] and summarized for use in the software POLYSOLVER [2] (see Methods).

| HLA-A Allele | Average Frequency |
|---|---|
| HLA-A*02:01 | 0.165735177970068 |
| HLA-A*24:02 | 0.115647218975036 |
| HLA-A*11:01 | 0.0988782090848313 |
| HLA-A*03:01 | 0.072759951608879 |
| HLA-A*01:01 | 0.0677237081068929 |
| HLA-A*23:01 | 0.0473178346593762 |
| HLA-A*33:03 | 0.0393454654639734 |
| HLA-A*30:01 | 0.0318933412012099 |
| HLA-A*26:01 | 0.031566894866778 |
| HLA-A*02:07 | 0.0269360269360269 |
| HLA-A*30:02 | 0.0238301274752413 |
| HLA-A*68:02 | 0.0238287779953814 |
| HLA-A*68:01 | 0.0231422454516778 |
| HLA-A*31:01 | 0.0228448436042297 |
| HLA-A*29:02 | 0.0214702245703172 |
| HLA-A*02:03 | 0.0175084175084175 |
| HLA-A*74:01 | 0.0174565392088051 |
| HLA-A*32:01 | 0.0171012988590301 |
| HLA-A*02:06 | 0.0164949325753346 |
| HLA-A*02:02 | 0.015105065441339 |
| HLA-A*25:01 | 0.0140723759785838 |
| HLA-A*34:02 | 0.0114125511748066 |
| HLA-A*02:05 | 0.0107374789865795 |
| HLA-A*33:01 | 0.0100718736040359 |
| HLA-A*66:01 | 0.00939507878432107 |
| HLA-A*36:01 | 0.00872776099362202 |
| HLA-A*11:02 | 0.00740740740740741 |
| HLA-A*24:07 | 0.003029285808742 |
| HLA-A*29:01 | 0.00302386744263803 |
| HLA-A*66:02 | 0.00234978180597516 |
| HLA-A*80:01 | 0.00234978180597516 |
| HLA-A*30:04 | 0.00201444117246695 |
| HLA-A*01:02 | 0.00201409869083585 |
| HLA-A*34:01 | 0.00168248446194066 |
| HLA-A*26:02 | 0.00134680134680135 |

| | |
|---|---|
| HLA-A*26:03 | 0.00134680134680135 |
| HLA-A*24:03 | 0.00134409216374936 |
| HLA-A*03:02 | 0.00134172546232848 |

**Supplementary Table S3:** Top frequency HLA-B alleles (n=81) used for neoepitope prediction from TCGA somatic mutations. Average allele frequencies in the general population are based on those frequencies originating from the Allele Frequency Net Database [1] and summarized for use in the software POLYSOLVER [2] (see Methods).

| **HLA-B Allele** | **Average Frequency** |
|---|---|
| HLA-B*07:02 | 0.0658965469288718 |
| HLA-B*15:01 | 0.0631742058525538 |
| HLA-B*40:01 | 0.0574267368967641 |
| HLA-B*35:01 | 0.0476382286835008 |
| HLA-B*08:01 | 0.0469771669978825 |
| HLA-B*53:01 | 0.0381638104006582 |
| HLA-B*51:01 | 0.0378425275710676 |
| HLA-B*44:03 | 0.0364888924510866 |
| HLA-B*18:01 | 0.0337932678573997 |
| HLA-B*58:01 | 0.0324242193820546 |
| HLA-B*40:02 | 0.0320892357921403 |
| HLA-B*44:02 | 0.0294062844017529 |
| HLA-B*46:01 | 0.027693346842283 |
| HLA-B*15:03 | 0.0226278332316052 |
| HLA-B*13:02 | 0.0223041527689579 |
| HLA-B*15:02 | 0.022289766970618 |
| HLA-B*45:01 | 0.0192516233688389 |
| HLA-B*42:01 | 0.0192502532928065 |
| HLA-B*52:01 | 0.0182411922949664 |
| HLA-B*27:05 | 0.0165628491553139 |
| HLA-B*58:02 | 0.016548463356974 |
| HLA-B*49:01 | 0.015202363655171 |
| HLA-B*13:01 | 0.0151975683890578 |
| HLA-B*57:01 | 0.0141957002903876 |
| HLA-B*14:02 | 0.0141909050242743 |
| HLA-B*39:01 | 0.0138514686872548 |
| HLA-B*57:03 | 0.0114829497462962 |
| HLA-B*35:03 | 0.0111544740175356 |
| HLA-B*38:01 | 0.0108170927945646 |
| HLA-B*15:10 | 0.00979398851739277 |
| HLA-B*54:01 | 0.00911854103343465 |
| HLA-B*37:01 | 0.00810982255460267 |
| HLA-B*56:01 | 0.00810913751658648 |
| HLA-B*15:18 | 0.00810605484551365 |
| HLA-B*38:02 | 0.00776764606551841 |
| HLA-B*48:01 | 0.00743026484254744 |

| | |
|---|---|
| HLA-B*55:02 | 0.00742992232353934 |
| HLA-B*81:01 | 0.00742992232353934 |
| HLA-B*50:01 | 0.00709630880965739 |
| HLA-B*55:01 | 0.0060834801027283 |
| HLA-B*41:02 | 0.0054070050617459 |
| HLA-B*07:05 | 0.0050665411677021 |
| HLA-B*40:06 | 0.00506585612968592 |
| HLA-B*14:01 | 0.00472984498274732 |
| HLA-B*15:25 | 0.00472813238770686 |
| HLA-B*27:02 | 0.004394861392833 |
| HLA-B*15:11 | 0.00439040864572779 |
| HLA-B*15:16 | 0.00439040864572779 |
| HLA-B*35:02 | 0.00405611009382966 |
| HLA-B*27:04 | 0.00405268490374873 |
| HLA-B*41:01 | 0.00371667375681013 |
| HLA-B*78:01 | 0.00371496116176967 |
| HLA-B*15:17 | 0.00304054123483583 |
| HLA-B*44:05 | 0.00202839756592292 |
| HLA-B*15:32 | 0.00202634245187437 |
| HLA-B*35:05 | 0.00202634245187437 |
| HLA-B*42:02 | 0.00202634245187437 |
| HLA-B*57:02 | 0.00202634245187437 |
| HLA-B*35:08 | 0.00169033130493577 |
| HLA-B*39:06 | 0.00168998878592768 |
| HLA-B*47:01 | 0.00168964626691958 |
| HLA-B*15:07 | 0.00168861870989531 |
| HLA-B*39:10 | 0.00168861870989531 |
| HLA-B*51:02 | 0.00168861870989531 |
| HLA-B*15:27 | 0.00135089496791624 |
| HLA-B*27:03 | 0.00135089496791624 |
| HLA-B*67:01 | 0.00135089496791624 |
| HLA-B*18:03 | 0.00101385626395337 |
| HLA-B*14:03 | 0.00101317122593718 |
| HLA-B*44:10 | 0.00101317122593718 |
| HLA-B*59:01 | 0.00101317122593718 |
| HLA-B*07:04 | 0.000676132521974307 |
| HLA-B*51:07 | 0.000675790002966215 |
| HLA-B*15:05 | 0.000675447483958122 |
| HLA-B*15:08 | 0.000675447483958122 |
| HLA-B*40:12 | 0.000675447483958122 |
| HLA-B*47:03 | 0.000675447483958122 |
| HLA-B*48:03 | 0.000675447483958122 |
| HLA-B*57:04 | 0.000675447483958122 |
| HLA-B*82:01 | 0.000675447483958122 |
| HLA-B*51:08 | 0.000338066260987153 |

**Supplementary Table S4:** Top frequency HLA-C alleles (n=26) used for neoepitope prediction from TCGA somatic mutations. Average allele frequencies in the general population are based on those frequencies originating from the Allele Frequency Net Database [1] and summarized for use in the software POLYSOLVER [2] (see Methods).

| HLA-C Allele | Average Frequency |
|---|---|
| HLA-C*07:02 | 0.116050870147256 |
| HLA-C*04:01 | 0.114899598393574 |
| HLA-C*07:01 | 0.0978888888888889 |
| HLA-C*06:02 | 0.0861914323962517 |
| HLA-C*03:04 | 0.0815890227576975 |
| HLA-C*01:02 | 0.0689317269076305 |
| HLA-C*02:02 | 0.0464230254350736 |
| HLA-C*03:03 | 0.0441619812583668 |
| HLA-C*16:01 | 0.0390281124497992 |
| HLA-C*12:03 | 0.0381298527443106 |
| HLA-C*08:01 | 0.0361419009370817 |
| HLA-C*05:01 | 0.0297510040160643 |
| HLA-C*17:01 | 0.0293507362784471 |
| HLA-C*03:02 | 0.0274163319946452 |
| HLA-C*08:02 | 0.0213668005354752 |
| HLA-C*12:02 | 0.0210816599732262 |
| HLA-C*14:02 | 0.0207336010709505 |
| HLA-C*15:02 | 0.0177309236947791 |
| HLA-C*07:04 | 0.0160441767068273 |
| HLA-C*18:01 | 0.01 |
| HLA-C*14:03 | 0.00769210174029451 |
| HLA-C*04:03 | 0.00602409638554217 |
| HLA-C*15:05 | 0.00600937081659973 |
| HLA-C*08:04 | 0.00333333333333333 |
| HLA-C*02:10 | 0.003 |
| HLA-C*08:03 | 0.00234270414993307 |

**Supplementary Table S6. Neoepitope sequence repetition within and across disease sites for real and simulated neoepitopes. Neoepitopes were randomly simulated for each disease site (see Methods), and the percentage of true versus randomly simulated neoepitope sequences repeated both within and across disease sites were compared. Significance of the difference in repetition between true and randomly simulated neoepitopes was tested using a Welch Two Sample t-test. For KICH, no epitope sequences were repeated within the disease site for either true or randomly simulated neoepitopes.**

| Disease Site (TCGA ID) | Mean % true epitopes repeated within site | Mean % random epitopes repeated within site | Significance of difference (p value) | Mean % true epitopes repeated across TCGA | Mean % random epitopes repeated across TCGA | Significance of difference (p value) |
|---|---|---|---|---|---|---|
| ACC | 0.38 | 0.027 | 0.0096 | 3.3 | 2.3 | 0.049 |
| BRCA | 4.5 | 0.34 | $< 2.2 \times 10^{-16}$ | 7.5 | 2.2 | $< 2.2 \times 10^{-16}$ |
| CESC | 2.5 | 0.025 | $< 2.2 \times 10^{-16}$ | 6.4 | 2.3 | $< 2.2 \times 10^{-16}$ |
| CHOL | 0.48 | 0.0016 | 0.0094 | 4.4 | 2.3 | 0.0024 |
| DLBC | 0.54 | 0.016 | 0.0044 | 4.2 | 2.3 | 0.0016 |
| KICH | 0 | 0 | NA | 4.9 | 2.4 | 0.0026 |
| KIRP | 0.41 | 0.083 | $1.2 \times 10^{-5}$ | 2.9 | 2.3 | 0.0059 |
| LAML | 12.4 | 0.026 | $2.4 \times 10^{-13}$ | 17.0 | 2.5 | $4.8 \times 10^{-14}$ |
| MESO | 2.1 | 0.013 | 0.0064 | 4.4 | 2.3 | 0.024 |
| PCPG | 2.0 | 0.0084 | 0.00032 | 6.4 | 2.3 | $6.9 \times 10^{-5}$ |
| PRAD | 1.9 | 0.11 | $< 2.2 \times 10^{-16}$ | 6.5 | 2.3 | $< 2.2 \times 10^{-16}$ |
| READ | 2.8 | 0.2 | $< 2.2 \times 10^{-16}$ | 7.1 | 2.3 | $< 2.2 \times 10^{-16}$ |
| SARC | 0.60 | 0.078 | $1.8 \times 10^{-6}$ | 4.4 | 2.3 | $3.2 \times 10^{-11}$ |
| SKCM | 12.1 | 1.2 | $< 2.2 \times 10^{-16}$ | 13.4 | 2.3 | $< 2.2 \times 10^{-16}$ |
| TGCT | 2.0 | 0.025 | 0.00042 | 4.5 | 2.4 | 0.0023 |
| THYM | 16.4 | 0.0043 | $< 2.2 \times 10^{-16}$ | 19.7 | 2.2 | $< 2.2 \times 10^{-16}$ |
| UCS | 2.2 | 0.022 | $2.9 \times 10^{-10}$ | 7.5 | 2.3 | $2.5 \times 10^{-9}$ |
| UVM | 14.9 | 0.0 | $< 2.2 \times 10^{-16}$ | 17.7 | 2.6 | $< 2.2 \times 10^{-16}$ |

**Supplementary Table S7:** Significance of difference in tumor vs. paired normal peptide sequence similarity for neoepitopes with non-anchor-position mutations with vs. without a putatively novel binding change. Across five neoepitope binding affinity windows, the significance of the difference in tumor vs. paired normal peptide sequence similarity between neoepitopes with and without a putatively novel binding affinity difference (see Methods) was tested with a Wilcoxon rank sum test.

| Neoepitope binding affinity (nM) | # neoepitopes with novel binding change | Median peptide sequence similarity for neoepitopes with novel binding change | # neoepitopes without novel binding change | Median peptide sequence similarity for neoepitopes without novel binding change | Significance of difference in peptide sequence similarity score between groups (p value) |
|---|---|---|---|---|---|
| < 100 | 140884 | 82.1% | 2098365 | 84.4% | $< 2.2\text{x}10^{-16}$ |
| > 100 and < 200 | 172779 | 82.4% | 1240385 | 84.6% | $< 2.2\text{x}10^{-16}$ |
| > 200 and < 300 | 126533 | 82.4% | 991259 | 84.6% | $< 2.2\text{x}10^{-16}$ |
| > 300 and < 400 | 101427 | 82.4% | 857704 | 84.6% | $< 2.2\text{x}10^{-16}$ |
| > 400 and < 500 | 85149 | 82.1% | 774308 | 84.6% | $< 2.2\text{x}10^{-16}$ |
